# Biotic interactions are more important at species’ warm vs. cool range-edges: a synthesis

**DOI:** 10.1101/2021.04.07.438721

**Authors:** Alexandra Paquette, Anna L. Hargreaves

## Abstract

Predicting which ecological factors constrain species distributions is a fundamental question in ecology and critical to forecasting geographic responses to global change. Darwin hypothesized that abiotic factors generally impose species’ high-latitude and high-elevation (typically cool) range limits, whereas biotic interactions more often impose species’ low-latitude/low-elevation (typically warm) limits, but empirical support has been mixed. Here, we clarify three predictions arising from Darwin’s hypothesis, and show that previously mixed support is partially due to researchers testing different predictions. Using a comprehensive literature review (886 range limits), we find that biotic interactions, including competition, predation, and parasitism, influenced species’ warm limits more often than species’ cool limits. At cool limits, abiotic factors were consistently more important than biotic interactions, but temperature contributed strongly to cool and warm limits. Our results suggest that most range limits will be sensitive to climate warming, but warm limit responses will depend strongly on biotic interactions.

> *“When we travel southward and see a species decreasing in numbers, we may feel sure that the cause lies quite as much in other species being favored, as in this one being hurt. (Whereas)… the number of species, and therefore of competitors, deceases northwards; hence in going northward or in ascending a mountain, we far oftener meet with stunted forms, due to the directly injurious action of climate”* –Darwin 1859

## INTRODUCTION

Most species distributions are constrained by poor fitness at and beyond range limits (Hargreaves et al. 2014, Lee-Yaw et al. 2016). Understanding and predicting which ecological factors limit fitness and therefore species distributions is a fundamental goal of ecology and biogeography (von Humboldt and Bonpland 1805, MacArthur 1984) and increasingly important for conservation. For example, knowing the extent to which temperature controls species distributions is critical to forecasting species range shifts under climate warming (Loarie et al. 2009, Sunday et al. 2015), whereas identifying natural enemies that govern species’ native ranges can inform biological control programs to limit their spread as invasive species (van Driesche et al. 2009). However, predicting which factors impose a given range limit requires a broad understanding of when abiotic vs. biotic constraints are most likely to limit fitness or dispersal, and such generalizations have remained elusive.

Studies of ecological constraints on species distributions often focus on abiotic constraints, likely because climate plays such a clear role in governing the current and historical distribution of terrestrial biomes (Emanuel et al. 1985, Castañeda et al. 2016). Abiotic variables like temperature and precipitation are easily quantified in standardized units, and climate data have been recorded for centuries (Bar-Yosef 2008, Slonosky 2014) and recently at a global scale (Yang et al. 2016). This makes climatic variables relatively easy to include in models, such that major platforms for modelling species distributions (e.g. BIOCLIM, Maxent) use climate as their major predictor (Phillips et al. 2006, Booth et al. 2014). However, climate-based distribution models can significantly mispredict species’ ranges (Bayly and Angert 2019), and species often fail to occupy regions with seemingly suitable climates (Früh et al. 2017) and occupy different abiotic niches on islands vs. mainland ecosystems (e.g. MacArthur 1984, Velasco et al. 2016). Thus abiotic factors are often insufficient to explain species distributions (Wisz et al. 2013).

Though less well-studied at range limits, biotic interactions can also limit species distributions. Biotic interactions affect fitness and structure communities (Mitchell et al. 2006, Wardle 2006), and these effects can be strong enough to limit a species’ range (Brown and Vellend 2014). However, quantifying biotic interactions at large scales is challenging since there is no common unit or tool for measuring interaction strength, and no significant history of recording it (Wiens 2011, Wisz et al. 2013). Given these difficulties it is unsurprising that few species distribution models include biotic interactions as predictors, yet we know we may be missing the full picture without them (Pearson and Dawson 2003, Wisz et al. 2013). To improve our understanding of ecological and biogeographic processes and target data collection to help predict species’ responses to global change, we need a predictive framework for when and where biotic interactions matter.

One such framework, first put forward by Darwin (1859), hypothesizes that the importance of biotic interactions varies predictably with latitude and elevation. Darwin, and others subsequently, suggested that toward the high-latitude and elevation edges of a species distribution (‘cool’ range limits), climate is the predominant factor that limits fitness and prevents range expansion, whereas at low-latitude and elevation edges (‘warm’ range limits), fitness is increasingly constrained by biotic interactions (Darwin 1859, MacArthur 1984). Increased importance of biotic interactions could arise if interactions become stronger in warmer, more productive, or more species-rich ecosystems (Darwin 1859, Roslin et al. 2017, Hargreaves et al. 2019), or simply because biotic interactions are relatively more important where abiotic conditions (especially climate) are more benign (Dobzhansky 1950). While Darwin’s conjecture that interactions are particularly important at warm range limits is well-known (Brown et al. 1996), it has proven tricky to test definitively.

The broad, general nature of Darwin’s conjecture is part of its appeal, but has led to disagreement about how it should be tested. One approach is to compare the importance of biotic interactions at contrasting range limits, with the prediction that biotic interactions will contribute more often or more strongly to warm vs. cool limits (*Prediction 1*; Fig. 1A). An alternate approach is to compare the relative importance of biotic vs abiotic factors at a given type of range limit. One might predict that at warm range limits, biotic factors will be more important than abiotic factors (*Prediction 2;* Fig. 1B), with the reverse true at cool limits. A third approach simply predicts that biotic interactions will contribute to a majority (>50%) of warm range limits (*Prediction 3*; Fig. 1C). All predictions could be true at once (Fig. 1D), but need not be. For example, biotic factors could be less important than abiotic factors everywhere but still be more important at warm vs. cool limits (i.e. support for *Prediction 1* but not *2*).

**Fig 1.**
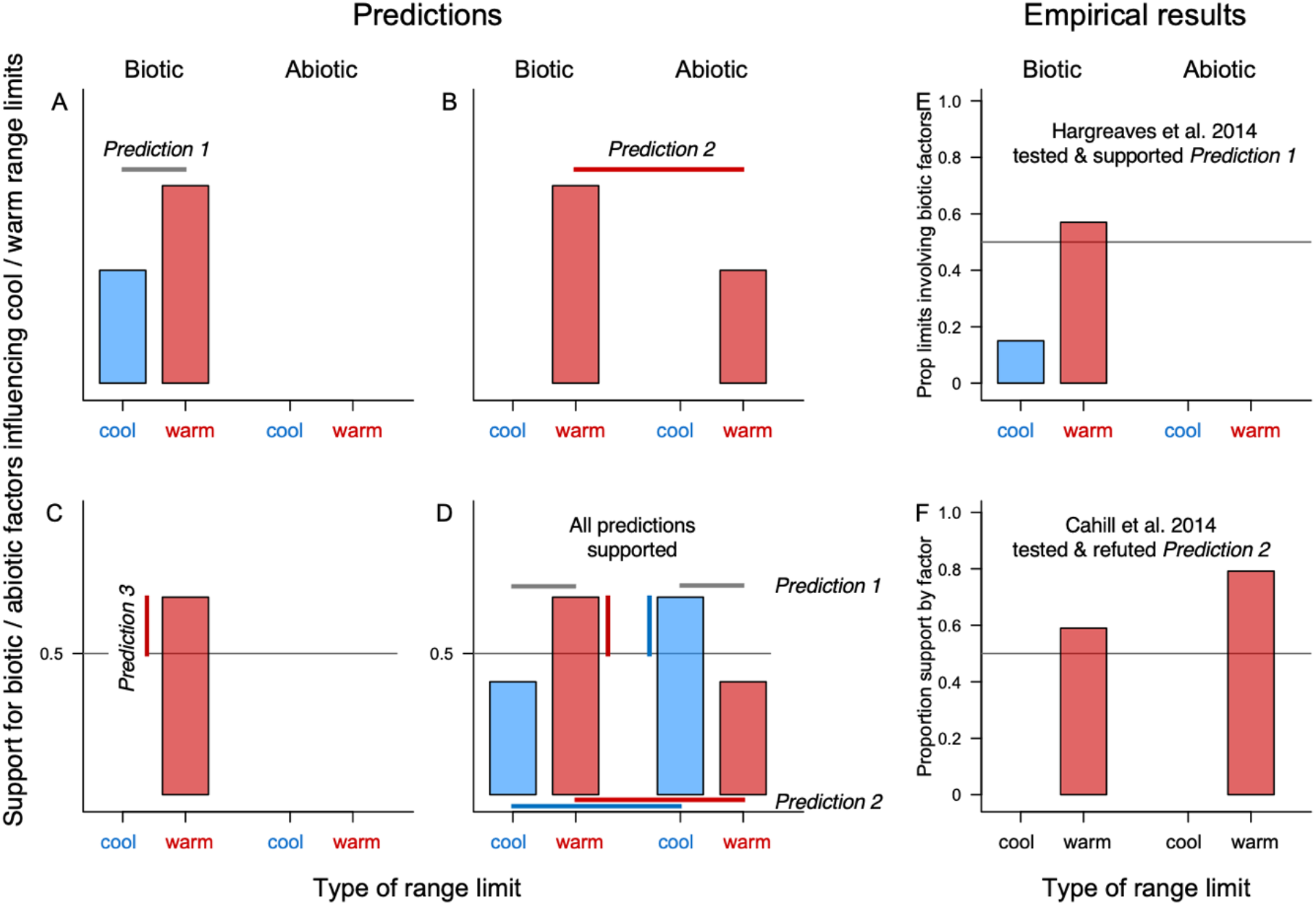
Assessing Darwin’s conjecture. A-D): Hypothetical results supporting predictions derived from Darwin’s conjecture: A) *Prediction 1*–biotic factors are more important at the warm (red) vs. cool (blue) limit of species ranges; B) *Prediction 2*–biotic factors are more important than abiotic factors at warm limits; C) *Prediction 3*–biotic factors contribute to >50% of warm limits; D) All three predictions could be true at once, along with their equivalent predictions for abiotic factors (i.e. abiotic factors are more important at cool vs. warm limits, more important than biotic factors at cool limits, important at >50% of cool limits). E-F): Results from past syntheses. E) Hargreaves et al. (2014) reviewed experiments that transplanted species beyond their warm or cool range limit. Response was whether biotic interactions contributed to the range limit, and results supported *Prediction 1* (and *Prediciton 3*). F) Cahill et al. (2014) reviewed field, lab, and modelling studies of factors contributing to warm range limits. Response was whether each factor assessed contributed to the range limit, and results refuted *Prediction 2* (but supported *Prediction 3*).

While several studies have tested Darwin’s conjecture for individual species (Merrill et al. 2008, Ettinger et al. 2011) or areas (e.g. Normand et al. 2009), two syntheses to date have tested its predictions globally; they tested different predictions, considered different data, and reached contrasting conlusions. Hargreaves et al. (2014) reviewed studies that transplanted species beyond their range and identified ecological factors that limited fitness beyond a warm or cool range limit. Hargreaves et al. tested and found strong support for *Prediction 1*; biotic factors contributed almost 4x more often to warm vs. cool limits (57% vs. 15%; Fig. 1E). In contrast, Cahill et al. (2014) reviewed field, lab, and modelling studies that assessed whether biotic or abiotic factors contributed to species warm range limits. Cahill et al. (2014) tested and refuted *Prediction 2*; abiotic factors contributed to warm range limits more often than biotic factors (79% of 164 tests vs. 59% of 49 tests; Fig. 1F). Thus Hargreaves et al. (2014) supported Darwin’s conjecture while Cahill et al. (2014) rejected it (neither study explicitly tested *Prediction 3*, but both supported it: Fig. 1), and it remains unclear whether the discrepancy is because they considered different data or tested different predictions.

Here, we test all three predictions derived from Darwin’s conjecture and resolve the apparently contradictory results of earlier reviews. We do this by synthesizing data on the drivers of warm and cool range limits from studies published up to 2019, and running three sets of analyses. First, we test the predictions in Fig 1, that biotic factors are: more important at warm vs. cool range limits (*Prediction 1*); more important than abiotic factors at warm limits (*Prediction 2*); and/or supported more often than not at warm limits (*Prediction 3*). Models simultaneously test the corresponding predictions for abiotic factors (supported more at cool vs. warm limits, more than biotic factors at cool limits, and more often than not at cool limits). We explore whether patterns are consistent between methods, latitudinal and elevational ranges, and terrestrial and marine environments. Second, we test which range-limiting factors are supported most often, as some have been predicted to be particularly important (e.g. temperature at cool limits, competition at warm limits; Darwin 1859, MacArthur 1984). Third, we test whether the relative support for biotic drivers increases toward the equator, as might be additionally expected if interactions are stronger toward the tropics (Dobzhansky 1950).

## METHODS

### Literature search

We searched Web of Science for studies published up to the end of 2019 that assessed the causes of species’ high-latitude or elevation (hereafter ‘cool’) or low-latitude or elevation (hereafter ‘warm’) range limits. We used a slightly modified version of the search terms in Cahill et al. (2014). The full term (terms we added are italicized) was: ((species (range or distribution) (border* or boundar* or edge* or limit* or margin*)) AND (biogeograph* or geograph* or global or altitud* or elevation* *or latitud**\****)) *AND (test* or experiment*)* AND (caus* or determin* *or driv**\*** or mechanis**\****). To increase coverage in areas with few studies, we repeated the search in Spanish and French and did a targeted search for studies from Africa (Supplementary Information 1). This yielded >2300 studies, which we then screened for those that assessed the importance of at least one biotic or abiotic factor in causing a cool or warm range limit. We added additional relevant studies from Hargreaves et al. (2014), Cahill et al. (2014), and our personal libraries. While Cahill et al. (2014) included some resurvey studies, which compare modern and historic ranges for a suite of species to test for range shifts expected under climate warming, we excluded resurveys as they have been reviewed extensively already (e.g. Freeman et al. 2018, Lenoir et al. 2020). We also excluded review papers to avoid duplicating results.

### Data extraction

We provide a detailed extraction protocol in SI.2, and summarize the key points here. We extracted data for each potentially range-limiting factor assessed, following Cahill et al. (2014) except that we separated data by study and species whenever possible (full details on how our data relate to Cahill et al. (2014)’s are in SI.3). For each study x taxon x range limit (latitude or elevation, cool or warm, separated by continent or ocean if applicable), we identified the potential range-limiting factors assessed, such that each factor a study assessed at a given range limit contributed 1 data point. We noted whether each factor was biotic or abiotic (‘factor type’) and what category of factor it was. Abiotic categories = temperature, precipitation/moisture, climate (when the effects of temperature and moisture could not be separated from each other or other climate variables), or soil (e.g. pH, nutrient content). Biotic categories = competition, biogenic habitat, predation/herbivory, host/food availability, or pathogens (i.e. parasitism or disease). Multiple assessments of a factor category (e.g. max. and mean annual temperature) at one species’ range limit would contribute one data point. Factors outside these categories were assigned as ‘abiotic other’ (e.g. salinity, light levels) or ‘biotic other’ (e.g. pollination), and not grouped (i.e. a study could contribute multiple data points for ‘abiotic other’).

We noted whether results came from field experiments (e.g. experimental manipulations of biotic or abiotic factors at the range edge), observational data (e.g. demographic surveys that correlate spatial variation in performance to abiotic or biotic conditions), lab experiments (mostly studies of thermal tolerance that compare tolerance limits to conditions at a species range limit), or models (e.g. species distribution models that evaluate whether large-scale environmental data predict the position of a species range limit); details in SI.2. If a study used multiple methods to reach a conclusion, we classified it under the method we considered most able to establish causation (field exp. > lab exp. > obs. > model); the latter three methods all rely on correlation to assess a factor’s importance at a given range limit. When possible, we recorded the latitude and longitude of each range limit (details in SI.4).

We assessed whether each factor contributed to the given range limit (‘yes’ or ‘no’), determined from statistical results, figures, and arguments in the discussion when necessary. Multiple factors could contribute to one range limit, e.g., the results of competitive interactions might depend on temperature (Davis et al. 1998). If a study considered multiple measures of one factor (e.g. summer and winter temperature), we deemed the factor (temperature in this example) supported if any measure contributed to the range limit. For studies in Cahill et al. (2014), we used their conclusions unless data were grouped across species or studies; in these cases we ungrouped data and reassessed conclusions for each species/study. As conclusions sometimes relied on integrating across results in a study, a second person spot-checked both our conclusions and those in Cahill et al. (2014); our archived data documents the reasoning and data behind each decision (SI.2).

### Data structure

We compared support for abiotic and biotic factors in two ways. First, following Cahill et al. (2014), we considered each factor assessed at one taxon x range limit as a binomial data point (yes if it contributed to the range limit). This ‘support by factor’ approach preserves the maximum power of the raw data. However, imagine a range edge imposed entirely by temperature. A study that tested the effect of temperature, precipitation, and soil pH (i.e. 3 data points for ‘abiotic’) would yield only 33% support for abiotic factors even though the range edge is 100% abiotically determined. We therefore also considered a second approach, ‘support by factor type’, which groups factors by type (abiotic or biotic), and asks ‘if at least one abiotic/biotic factor was tested at a study x taxon x range limit, was at least one abiotic/biotic factor supported?’. Thus, the example above would contribute 1 data point for ‘abiotic’ (‘yes’ since temperature was supported), and none for ‘biotic’ since no biotic drivers were tested. We included the number of factors tested per factor type (n = 3 for abiotic factors in the example above) as a covariate; this was particularly important as studies tended to assess fewer biotic factors per taxon x range limit (max. 3) than abiotic factors (max. 5). Results were consistent between the two response variables, so we present the ‘support by factor’ results here and give the ‘support by factor type’ results for Analysis 1 in Table S1.

### Analyses

Analyses were done in R 3.4.2 (R Core Team 2017). Data and code will be publicly archived on publication. All analyses used binomial generalized linear mixed models (GLMMs) with ‘support by factor’ (i.e. 1 data point per factor assessed per taxon per range limit per study) as the response, logit link functions (lme4 package, v 1.1.23; Bates et al. 2015), and ‘species’ and ‘study’ as random factors. The significance of interaction terms was assessed using likelihood ratio tests compared to a Chi-squared distribution (*anova*, base R). To visualize results, we extracted back-transformed least-squared means, trendlines, and partial residuals from final models (using *visreg* and *emmeans* packages; (v 2.7.0 Breheny and Burchett 2017, v 1.5.1 Lenth et al. 2020)). The gap between partial residuals and predicted means in our figures indicates that the remaining model terms never exactly predicted the observed data.

#### Analysis 1)

To test our main predictions regarding the relative importance of biotic vs. abiotic factors at warm limits (Fig. 1), we ran a GLMM with ‘factor type’ (biotic or abiotic), ‘range limit type’ (cool or warm), and their interaction as fixed effects. The factor type x range limit type interaction was significant so was retained in the final model. To assess whether support differed between cool and warm range limits for a given factor type (*Prediction 1*) or between biotic and abiotic factors at a given range limit type (*Prediction 1*), we computed least-squares means and post-hoc comparisons among means. We determined that a given factor type was supported at a given range limit more often than not (*Prediction 3*) if 95% CI did not overlap with 0.5.

We explored the sensitivity and biological variation in our main results. To test sensitivity, we reran the model above on restricted data sets: i) including only field experiments, which are best able to establish causation; ii) excluding results grouped across multiple species; iii) including only taxon range limits where both biotic and abiotic factors were assessed; and iv) including only taxon ranges (latitudinal or elevational) where both cool and warm limits were assessed. We tested whether results varied between latitudinal and elevational ranges by rerunning our main and field-experiments-only analyses with an additional fixed effect ‘geographic range type’ (latitude or elevation), and assessing the importance of the 3-way interaction: factor type (biotic/abiotic) x range limit type (cool/warm) x geographic range type. Similarly, we tested whether results varied among terrestrial and marine environments, as biotic interaction strength seems to vary predictably with latitude on land (Roslin et al. 2017, Hargreaves et al. 2019) but not in oceans (Roesti et al. 2020). There were too few tests of freshwater systems to include them, and too few marine field experiments to robustly test the effect of environment for field experiments only (Table S2), so we reran our main analyses with a third interacting fixed effect (‘environment’, i.e. terrestrial or marine) and assessed the importance of the 3-way interaction.

#### Analysis 2)

We tested which categories of abiotic and biotic factors most often contributed to cool vs. warm range limits. We used the nine most common categories (Abiotic = temperature, precipitation/moisture, climate, soil; Biotic = competition, biogenic habitat, predation/herbivory, host/food availability, parasitism/disease); and excluded factors categorized as ‘other’. We tested whether support for each factor category differed among cool and warm limits using a GLMM with ‘factor category’, ‘range limit type’ (cool or warm), and their interaction as fixed effects. Importance of the interaction and differences among factor levels were assessed using a likelihood ratio tests and least-squared mean contrasts, as in Analysis 1.

#### Analysis 3)

We tested whether the conclusions to *Predictions 1* and *2* varied with latitude by rerunning our main model from Analysis 1 including ‘absolute latitude’ as a third fixed effect (factor type x range limit type x absLatitude). As the 3-way interaction was significant, we used post-hoc contrasts to test for differences in latitudinal trends for abiotic vs biotic factors at cool vs. warm range limits. For plotting, confidence intervals were estimated by bootstrapping the model 1000x following Bolker (2015). To test the sensitivity of results, we also ran this analysis for field experiments only, and separately for terrestrial and marine environments (SI.5).

## RESULTS

We identified 338 studies including 1941 assessments of potentially range-limiting factors. Data came from 656 taxa: 632 species or subspecies plus 28 groups for which results could not be separated by species (e.g. *Nebria* spp, ‘boreal trees’; Table 1). Most taxa were plants/algae (361, of which 86% were vascular land plants) or animals (292, of which 54% were vertebrates). Cool limits were studied more often than warm limits (Fig. S1) and abiotic factors were assessed more than twice as often as biotic factors (72% vs. 28% of data, respectively; Table 1). Data came from all seven continents but were dominated by temperate latitudes in the northern hemisphere (Fig. S1). Data were largely from terrestrial species (78% terrestrial, 19% marine, 3% freshwater; Table S2) but split fairly evenly between latitudinal and elevational limits (Table S3). Field experiments, which have the greatest power to establish causation, were well represented among studies (used in 36% of studies), but tended to test fewer factors than models and observational studies, so contributed only 28% of factor assessments (Table S2).

**Table 1.** Sample sizes. ‘Taxa’ are mostly individual species but include some species groups (e.g. heathland plants). Each ‘Range limit’ represents one taxon at either the cool or warm limit of either it’s latitudinal or elevational range on one continent or ocean (some range limits were studied in >1 study). Last two rows show the distribution of factor assessments, where each factor assessed (e.g. temperature, precipitation, competition) contributes a data point.

| Literature search results | Total | Cool limits | Warm limits |
| --- | --- | --- | --- |
| Studies <sup>a</sup> | 338 | 247 | 206 |
| Taxa <sup>b</sup> | 656 | 467 | 373 |
| Range limits (RL) | 886 | 491 | 395 |
| Study x taxon x RL x abiotic factor | 1407 | 771 | 636 |
| Study x taxon x RL x biotic factor | 534 | 289 | 245 |

a) Total is less than Cool + Warm as some studies assessed both cool and warm range limits
b) Total is less than Cool + Warm as some taxa were studied at both cool and warm limits

### 1) Relative importance of biotic vs. abiotic factors at cool vs. warm range limits

The relative importance of biotic and abiotic factors differed between range limits, supporting most predictions derived from Darwin’s conjecture (Fig. 1). Biotic factors were more important at warm range limits than at cool limits (supporting *Prediction 1*), and contributed to warm limits more than 50% of the times they were tested (supporting *Prediction 3*; Fig. 2). This was true across the data and in field experiments only (Fig. 2), and in all but one sensitivity test (Table 2). Abiotic factors were consistently more important than biotic ones at cool range limits (supporting *Prediction 2*), and contributed to cool limits more than 50% of the times they were tested (supporting *Prediction 3*; Fig 2, Table 2). Predictions that abiotic factors were more important at cool vs. warm limits (*Prediction 1*) and that biotic factors were more important than abiotic ones at warm limits (*Prediction* 2) received mixed support, being supported across data and in field experiments, respectively, and in some but not all sensitivity tests (Fig. 2; Table 2). The results above were consistent between latitudinal and elevational range limits (factor type x range limit type x geographic range type interaction NS), and between terrestrial and marine environments (factor type x range limit type x environment interaction NS; Table S3).

**Fig 2.**
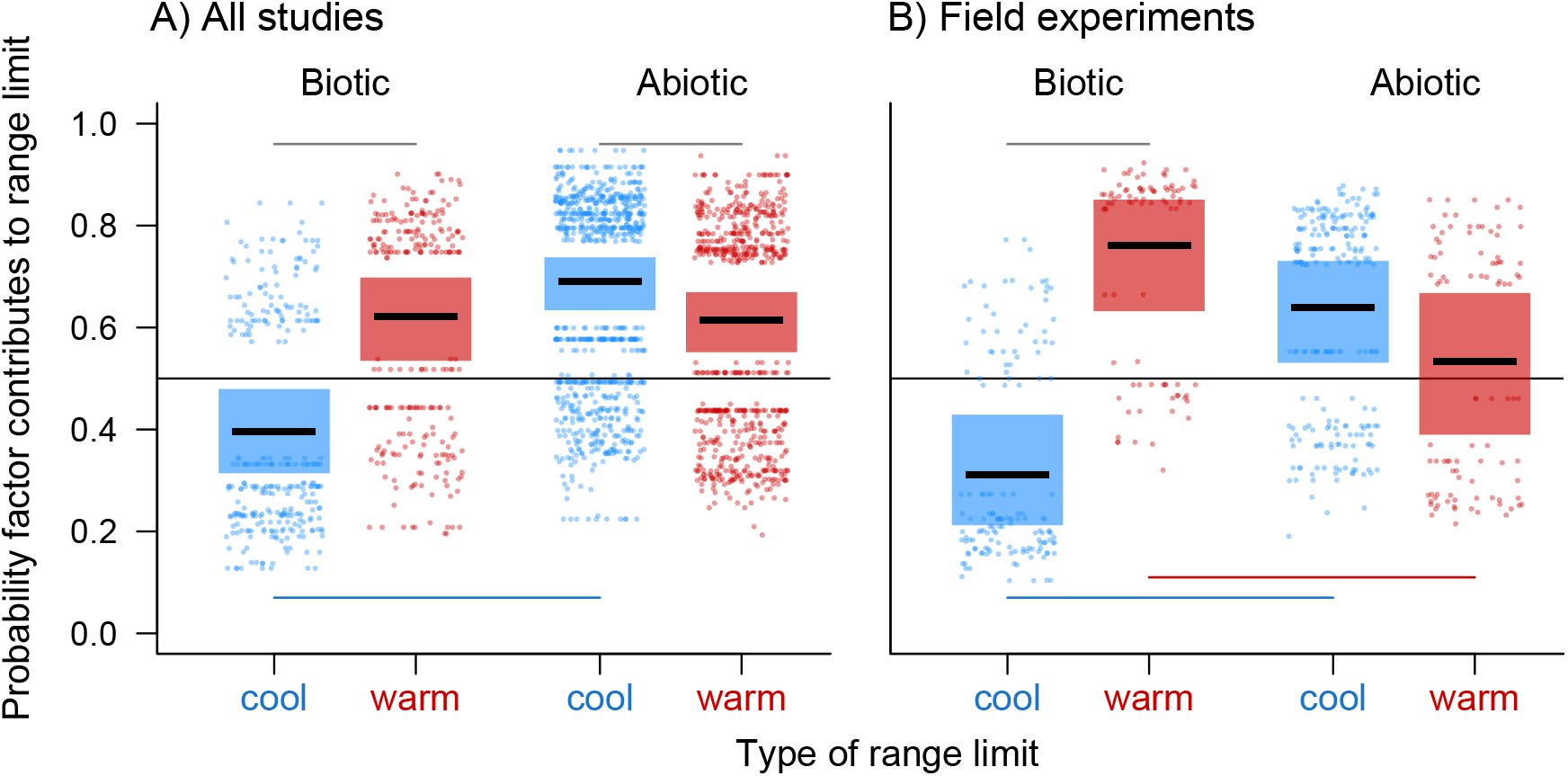
Importance of biotic and abiotic factors differs between cool and warm range limits. (A) All study types (field experiments, lab experiments, observational studies, models); (B) Experiments in nature. Top and bottom lines indicate significant pairwise differences supporting one of Darwin’s predictions. Top lines: biotic factors were more often important at warm vs cool limits, and abiotic factors were more often important at cool vs. warm limits (*Prediction 1*). Bottom lines: abioitc factors were more important than biotic ones at cool limits (blue); biotic factors were more important than abiotic ones at warm limits in field experiments (red; *Prediction 2*). In both panels, biotic and abiotic factors were supported >50% of the time at warm and cool limits, respectively (*Prediction 3*). Centre lines, boxes and points show means, 95% CI, and partial residuals extracted from final binomial GLMMs (horizontal variation in residuals is added for visualization).

**Table 2.** Statistical results for main analyses. We considered two response variables: support by factor (each factor assessed contributes one data point), and support by factor type (factors grouped as abiotic or biotic). Main analyses consider all data. Sensitivity analyses assess whether results are consistent for subsets of data that might be considered particularly powerful for testing the relative importance of biotic and abiotic factors in causing species’ cool vs. warm range limits. Predictions detailed in Fig. 1. * indicates results shown in Fig. 2.

| Data | n<br>total | factor type x<br>range limit type |  | *Prediction 1* |  | *Prediction 2* |  | *Prediction 3* |  |
| --- | --- | --- | --- | --- | --- | --- | --- | --- | --- |
|  |  | Chisq,<br>df=1 | *P* | Warm > Cool<br>for biotic | Cool > Warm<br>for abiotic | Bio > Abio<br>at warm | Abio > Bio<br>at cool | Bio >0.5<br>at warm | Abio >0.5<br>at cool |
| All* | 1941 | 24.0 | <0.001 | Y (<0.001) | Y (0.015) | N (0.87) | Y (<0.001) | Y | Y |
| Sensitivity analyses |  |  |  |  |  |  |  |  |  |
| i) Field experiments only* |  |  |  |  |  |  |  |  |  |
|  | 546 | 26.8 | <0.001 | Y (<0.001) | N (0.19) | Y (0.005) | Y (<0.001) | Y | Y |
| ii) Species-level data only |  |  |  |  |  |  |  |  |  |
|  | 1866 | 23.6 | <0.001 | Y (<0.001) | Y (0.013) | N (0.88) | Y (<0.001) | Y | Y |
| iii) Taxon range limits where both abiotic & biotic factors were assessed |  |  |  |  |  |  |  |  |  |
|  | 1118 | 8.6 | 0.003 | Y (<0.001) | N (0.96) | N (0.45) | Y (<0.001) | Y | Y |
| iv) Taxon elevational or latitudinal ranges where both cool & warm limits were assessed |  |  |  |  |  |  |  |  |  |
|  | 928 | 11.5 | <0.001 | N (0.054) | Y (<0.001) | N (0.23) | Y (<0.001) | N (=0.5) | Y |

### 2) Which factors contribute most often to cool and warm range limits

Different individual factors contributed most often to cool vs. warm range limits (Fig. 3). At cool limits, temperature contributed most often, and was the only factor supported more often than expected by chance (>50% of assessments) both across studies and in field experiments (Fig. 3A&B). Temperature and climate (which includes temperature effects) were supported more often than predation/herbivory and pathogens, whereas biotic factors were never supported more often than abiotic ones (Fig. 3A&B). At warm limits, biotic factors contributed most often, particularly predation/herbivory and competition which contributed to warm limits significantly more often than cool limits both across studies and in field experiments (Fig. 3C&D). Availability of appropriate biogenic habitat and moisture also frequently contributed to warm limits. Results from field experiments were similar to those found across studies for cool limits (Fig. 3 B vs A), but at warm limits field experiments found stronger support for competition and moisture, and weaker support for temperature (Fig. 3D vs C) compared to results across methods.

**Figure 3.**
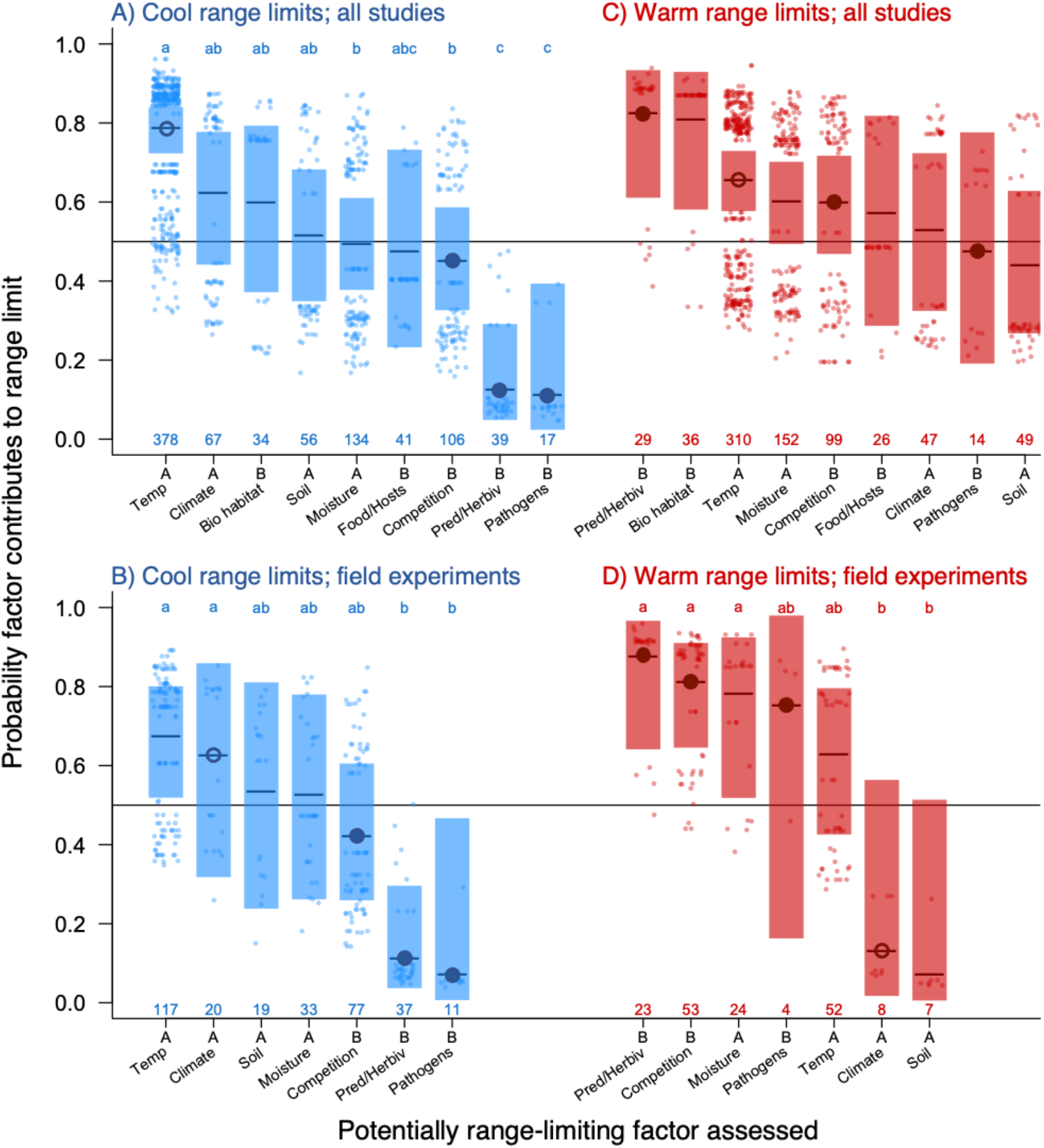
Different factors drive cool vs. warm range limits. The relative importance of the most commonly-assessed factors differed between cool and warm limits (factor x limit type: χ^2^df=8 = 50.9, *P* < 0.001 all studies (A&C); χ^2^df=6 = 44.4, *P* < 0.001 field experiments (B&D). Factors: A=abiotic, B=biotic, abbreivated names are: temperature, biogenic habitat, precipitation/moisture, food/host availability, predation/herbivory. Field experiments seldom assessed the effects of biogenic habitat or food/host availability, so analyses in B&D exclude these factors. Centre circles indicate factors whose importance differs between cool and warm limits (open = abiotic, closed = biotic). Top letters indicate significant differences among factors at a given limit (no factor pairs differed significantly at warm limits in A). Bottom numbers indicate the times the factor was assessed (study x taxon x range limit type combinations). Plots show back-transformed means, 95% CI and partial residuals extracted from binomial GLMMs.

### 3) Does the relative importance of biotic vs. abiotic factors vary with latitude?

Our results varied somewhat with latitude, but not as predicted. We predicted that biotic factors might become less important and abiotic factors more important toward the poles (i.e. factor type x absolute latitude interaction), but found that this varied between cool and warm range limits (i.e. factor type x range limit type x absolute latitude interaction: χ^2^_df=1_ = 4.5, *P* = 0.033; full statistical details in Table S4). Biotic factors became slightly less important and abiotic factors slightly more important toward the poles at cool range limits (as predicted; Fig. 4 blue lines), but showed opposite (biotic) or no (abiotic) trends at warm limits (Fig. 4 red lines), and none of the individual latitudinal trends were significant. Latitudinal trends did not differ by range limit type for either biotic or abiotic factors, nor between factor type at either cool or warm range limits, across systems (Fig. 4) or in land or marine environments (Fig. S2). Field experiments yielded similar results for biotic factors, and even less expected results for abiotic factors, whose importance *declined* toward the poles at warm limits (opposite of predictions; Fig. S2).

**Fig 4.**
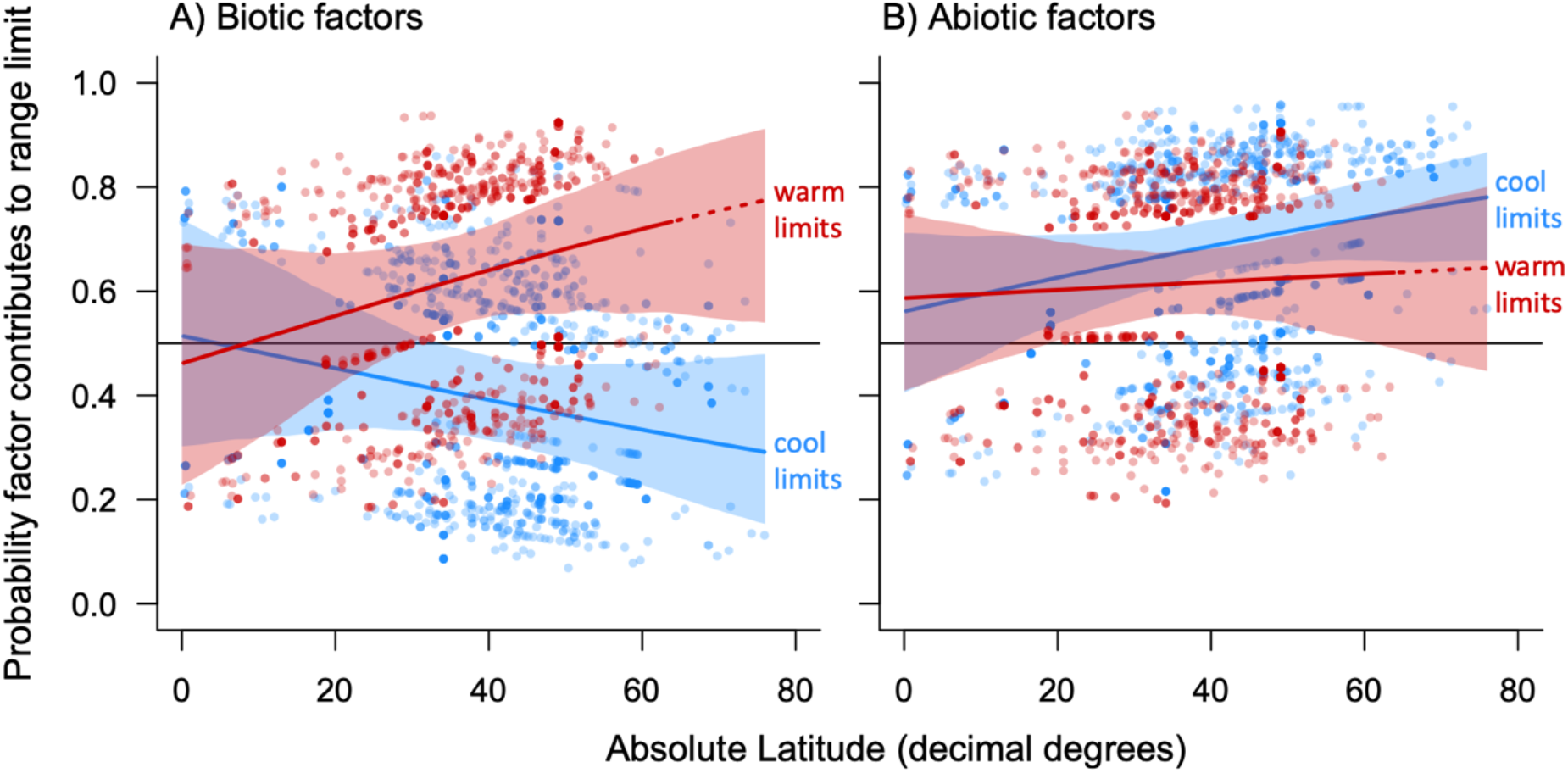
Latitudinal patterns in the importance of biotic and abiotic factors at range limits. The relative importance of biotic vs. abiotic factors at warm vs. cool range limits varied with distance from the equator (significant factor type x range limit type x absolute latitude interaction). Nevertheless, latitudinal trends did not vary significantly between range limit type for either biotic (A) or abiotic (B) factors (though the difference was almost significant in A, *P* = 0.06), or between cool (blue) and warm (red) range limits for either factor type (though the difference was almost significant at cool limits, *P* = 0.06; full statistical details in Table S4). None of the four latitudinal trends was significant. Lines, polygons and points show trend lines, 95% CI and partial residuals extracted from the full GLMM. Dashed lines extend predictions above latitudes where we had warm limit data (cool data extend to the equator due to high-elevation limits)

## DISCUSSION

Our review of >300 studies of the ecological limits to species’ ranges robustly supported some predictions derived from Darwin’s conjecture, and did not strongly refute any. Biotic interactions were consistently more important at species’ equatorward and low-elevation range edges than at their poleward and high-elevation range edges (Table 1). This was the prediction tested and supported by Hargreaves et al. (2014) using field experiments (2014; Fig. 1); we find it to be true both in field experiments and across study methods (Fig. 2). Second, abiotic factors were more important than biotic interactions at species’ cool range limits. This prediction has not, to our knowledge, been systematically tested before. Third, most warm limits were at least partially imposed by biotic interactions, and most cool limits were at least partially imposed by abiotic factors (Fig. 1; Table S1). Our results help resolve conflicting conclusions from past studies, and can inform both our understanding of ecology and efforts to predict future range shifts.

We show that previous conflicting conclusions about the importance of biotic interactions at warm range limits (Fig. 1) arise from testing different predictions and considering different data. Biotic interactions were more important at warm limits than at cool limits (*Prediction 1*; Fig. 2), and contributed to a majority of warm limits (*Prediction 3;* Table S1) no matter how they were assessed (Fig. 2). However, they were only found to be more important than abiotic factors at warm limits (*Prediction 2*) in field experiments, not across study methods. Varying support among predictions shows it is critical for studies to clearly articulate which prediction(s) they are testing, and acknowledge that refuting (or supporting) one does not negate or confirm Darwin’s conjecture as a whole.

Some interactions have particularly influenced ecological thinking about the asymmetric importance of biotic interactions. Both Darwin (1859) and MacArthur (1984) focused on competitive exclusion, and competition remains the most empirically assessed interaction at range edges (Fig. 3). Our synthesis found that competition contributed to most warm limits and to warm limits more than cool limits, a strikingly clear result given that competition was often inferred from proxies like relative growth rate or niche breadth (Cadena and Loiselle 2007). Similarly, pathogens shaped ideas that interactions play stronger ecological roles in tropical communities (Janzen 1970, Connell 1971, Schemske et al. 2009), and our results confirmed that pathogens constrain warm range limits more than cool ones despite remarkably few assessments. While our results support the theoretical importance of competitors and pathogens, predation/herbivory and the availability of biogenic habitat were equally or more important (Fig. 3). Predation has been the primary focus of large-scale tests of latitudinal gradients in interaction strength, which have generally found higher consumption at low latitudes and elevations (Jeanne 1979, Roslin et al. 2017, Hargreaves et al. 2019), consistent with a greater ecological role at warm range limits. To our knowledge biogenic habitat is rarely considered in discussions of Darwin’s conjecture; its strong contribution to warm limits suggests we may have been neglecting an important interaction.

Efforts to understand the ecological limits to species distributions have been reinvigorated by the desire to forecast range shifts under climate change. Our results support the long-held views that temperature influences most range edges, and that temperature is particularly important at cool range limits (Fig. 3). Whether this means it will be easier to predict expansions at cool limits than changes to warm range limits remains to be seen. First, it is unclear whether the support for temperature and precipitation in our data reflect a direct role of climate; climatic factors are tested far more than others, no doubt due to the availability of data, and are highly correlated with other potential drivers (Hargreaves et al. 2019). Indeed, field experiments find lower support for temperature, suggesting its effects on ranges may often be indirect. Predictive ability may also break down if climate change produces non-analogue climates (Ellis et al. 2017). Second, although biotic interactions were more important at warm limits, biogenic habitat, competition and food/host availability still influenced many cool limits (Fig. 3). As our analyses addresses only how often interactions contribute, not how strongly, it remains unclear whether predictions without interactions will be sufficient. To date, changes in species warm range limits have not been more varied than shifts in paired cool limits (Freeman et al. 2018). Both have moved poleward and upward on average, reflecting climate’s importance as seen in our data, but with high variability, indicating that non-climate factors may hamper fine-scale predictions.

In addition to revealing ecological patterns, syntheses reveal research biases, and we found considerable variation in how often ecological factors are assessed at range limits. Biotic factors were assessed far less often than abiotic factors (Table 1). This was particularly so in lab experiments and models, no doubt because these approaches often rely on large scale databases, which do not exist for interactions (except perhaps biogenic habitat). Pathogens are particularly poorly studied despite their prominence in theory. Mutualisms are notably absent from Fig. 4, and were only tested in 13 of 337 studies. Thus, as is often the case (e.g. Hargreaves et al. 2020), we have much better data about the influence of negative interactions than positive ones, despite evidence that lack of mutualist partners can dramatically affect range edges (Afkhami et al. 2014, Pither et al. 2018).

Our synthesis also revealed contrasting conclusions among methods, highlighting the importance of not just which data are collected but how. Field experiments offer an important advantage over approaches that rely on correlations to establish causation (Colwell and Rangel 2009), as multiple abiotic and biotic factors often covary spatially. Particularly powerful are experiments that manipulate putatively range-limiting factors in range-edge populations or beyond-range transplants (e.g. Griffith and Watson 2006, Hargreaves and Eckert 2019, Anderson and Wadgymar 2020). However, field experiments are often less geographically extensive and shorter than ideal for testing long-term constraints to distributions (Hargreaves et al. 2014). Most powerful are studies that combine the strengths of multiple methods, e.g. using experiments to establish a causal link and models or observational data to confirm patterns across larger spatial or temporal scales (e.g. Battisti et al. 2005, Afkhami et al. 2014). Because of this increased power, multi-method studies are best able to illuminate underappreciated causes of range limits (e.g. mutualisms; Afkhami et al. 2014); more of them would deepen our understanding of the ecological forces shaping species distributions.

How should range-limit research proceed from here? 1) Our understanding of species distributions would benefit greatly from more assessments of how biotic factors influence them. It is hard to comment definitively on the role of interactions at species range limits if we continue to assess them a third as often as we assess abiotic factors. 2) We re-iterate previous calls for more definitive tests of proximate causes of species range edges, particularly studies that assess paired warm and cool limits or that directly test the effects of biotic and abiotic factors. Combined tests of biotic and climatic factors would help determine their relative importance and how accurate climate-based range shift projections are likely to be. 3) Models suffer from the lack of widespread data on interactions, so finding reliable proxies would immensely beneficial. There are increasing large-scale tests for geographic patterns in the strength of interactions; these could be paired with assessments of potential drivers (e.g. productivity or species richness), to identify useful correlates for interaction strength, if they exist.

## ACKNOWLEDGEMENTS

We thank Ella Martin for help double-checking extracted data, and Jennifer Sunday and the Hargreaves and Sunday labs (McGill Biology) for thoughtful discussion about these ideas. Funding was provided by the Natural Science and Engineering Research Council of Canada via two Undergraduate Student Research Awards to AP and a Discovery Grant to ALH.

## Notes

### Competing Interest Statement

The authors have declared no competing interest.

